# Integrating reward information for prospective behaviour

**DOI:** 10.1101/2021.03.30.437719

**Authors:** Sam Hall-McMaster, Mark G. Stokes, Nicholas E. Myers

## Abstract

Value-based decision-making is often studied in a static context, where participants decide which option to select from those currently available. However, everyday life often involves an additional dimension: deciding *when* to select to maximise reward. Recent evidence suggests that agents track the latent reward of an option, updating changes in their latent reward estimate, to achieve appropriate selection timing (*latent reward tracking*). However, this strategy can be difficult to distinguish from one in which the optimal selection time is estimated in advance, allowing an agent to wait a pre-determined amount of time before selecting, without needing to monitor an option’s latent reward (*distance-to-goal tracking*). Here we show that these strategies can in principle be dissociated. Human brain activity was recorded using electroencephalography (EEG) while female and male participants performed a novel decision task. Participants were shown an option and decided when to select it, as its latent reward changed from trial-to-trial. While the latent reward was uncued, it could be estimated using cued information about the option’s starting value and value growth rate. We then used representational similarity analysis to assess whether EEG signals more closely resembled latent reward tracking or distance-to-goal tracking. This approach successfully dissociated the strategies in this task. Starting value and growth rate were translated into a distance-to-goal signal, far in advance of selecting the option. Latent reward could not be independently decoded. These results demonstrate the feasibility of using high temporal resolution neural recordings to identify internally computed decision variables in the human brain.

**Significance Statement:** Reward-seeking behaviour involves acting at the right time. However, the external world does not always tell us when an action is most rewarding, necessitating internal representations that guide action timing. Specifying this internal neural representation is challenging because it might stem from a variety of strategies, many of which make similar predictions about brain activity. This study used a novel approach to test whether alternative strategies could be dissociated in principle. Using representational similarity analysis, we were able to distinguish between candidate internal representations for selection timing. This shows how pattern analysis methods can be used to measure latent decision information in non-invasive neural data.

## Introduction

Neural research on decision-making has provided detailed accounts of how rewards are compared between options (A vs. B), to determine which option should be selected (e.g. Fouragnan et al., 2019; Hunt & Hayden, 2017; Murray & Rudebeck, 2018; Padoa-Schioppa, 2011; Rich & Wallis, 2016). However, everyday life also requires us to decide when to act on our selection (Khalighinejad et al., 2020a, 2020b). For example, we might have purchased an avocado to use during the week, but want to time when we use it, to ensure it is not under or overripe. Such a decision can be challenging to optimise because the rewards associated with an option are rarely stationary and, as a result, the timing of selection is critical to achieve the desired outcome.

Prior research has shown that the basal forebrain integrates recent experience with ongoing sensory information about an option’s reward prospect, to guide decisions about selection timing (Khalighinejad et al., 2020a, 2020b). But how such timing is calibrated without ongoing sensory reward cues remains largely unknown. Recent results indicate that the dorsal anterior cingulate cortex can encode latent information about an option’s progress to a reward threshold, in the absence of sensory input (Stoll et al., 2016). This suggests that option selection could be timed using an internal representation that continuously updates an estimate of the option’s latent reward (*latent reward tracking*). While this proposal holds theoretical appeal, optimal selection timing could also be achieved using prospection (e.g. Bulley et al., 2016) and delay estimation (e.g. Coull et al., 2011; Hassall et al., 2020), to anticipate when an option will be most rewarding in future and initiate selection when that time point is reached (*distance-to-goal tracking*).

Here we sought to develop an approach that could distinguish between these strategies. This is challenging because latent reward and distance-to-goal are often correlated; as reward increases towards a selection threshold, the distance to the selection event decreases. Therefore, even neural activity that appears to track one variable might be equally well explained by the other. Identifying which of these variables better fits neural activity is a critical step towards a mechanistic explanation of how the brain computes decision timing (O’Reilly & Mars, 2011). The present experiment recorded electroencephalograms (EEG) while human participants made selection timing decisions. Participants were shown an option that had a distinct starting value and growth rate. With each successive trial, the reward for deciding to select updated based on the growth rate, without corresponding sensory changes to indicate the current value. Pattern classification was then used to confirm that both an option’s latent reward and distance-to-goal could be decoded from the EEG signal, consistent with the fact that the two variables were correlated and, therefore, that the neural representation of either variable could explain the decoding in both cases. Representational similarity analysis (RSA) was then used to examine whether EEG signals corresponded more closely to latent reward tracking or distance-to-goal tracking, while controlling for each variable. We predicted that neural signals would be more consistent with latent reward tracking, based on prior indications that reward monitoring is used to initiate select decisions (Khalighinejad et al., 2020a, 2020b; Stoll et al., 2016),

The results from this experiment indicate that the two decision strategies could be dissociated. Neural responses were better explained by distance-to-goal tracking. Independent evidence for latent reward tracking was not detected. These results suggest that, in some contexts, decisions about when to select an option might be made by transforming option information into an estimate of optimal selection timing. At a broader level, we highlight the utility of time-resolved EEG decoding to disambiguate latent decision variables in human participants.

## Materials and Methods

### Participants

We set a target sample size of 32 participants based on previous M/EEG studies in our group (e.g. Hall-McMaster et al., 2019; Tankelevitch et al., 2020; Wolff et al., 2020). Five participants were excluded from the initial sample. One participant was excluded based on low behavioural performance, which fell more than three standard deviations below the mean percentage of points gained on the task (89.29%). Four participants were excluded based on excessive EEG artefacts, which lead to the rejection of more than 400 trials during pre-processing (> 21.27-27.47% of trials depending on the participant). To meet the 32 participant target, we therefore collected data from five additional participants. The final sample were between 18 and 35 (mean age=25, 19 female). All participants reported normal or corrected-to-normal vision (including normal colour vision) and no history of neurological or psychiatric illness. Participants received £10 per hour or course credit for taking part and could earn up to £20 extra for task performance. The study was approved by the Central University Research Ethics Committee at the University of Oxford (R58489/RE001) and all participants signed informed consent before taking part.

### Materials

Stimuli were presented on a 24-inch screen with a spatial resolution of 1920 x 1080 and refresh rate of 100Hz. Stimulus presentation was controlled using Psychophysics Toolbox-3 (RRID: SCR_002881) in MATLAB (RRID: SCR_001622, version R2015b). F and J keys on a standard QWERTY keyboard were used to record left and right hand responses. EEG data were recorded with 61 Ag/AgCl sintered electrodes (EasyCap, Herrsching, Germany), a NeuroScan SynAmps RT amplifier, and Curry 7 acquisition software (RRID: SCR_009546). Electrode impedance was reduced to <10 kΩ before recording. EEG data were pre-processed in EEGLAB (RRID: SCR_007292, version 14.1.1b) (Delorme & Makeig, 2004). Data were analysed in MATLAB (RRID: SCR_001622, version R2020a). Bonferroni-Holm corrections were implemented using David Groppe’s MATLAB toolbox (https://bit.ly/3xLBJlW). Bayesian paired t-tested were implemented using Bart Krekelberg’s MATLAB toolbox (https://klabhub.github.io/bayesFactor/). Bonus items used to incentivise performance were created by icon developers Smashicons, Freepik, Flat Icons, Goodware, Pixel Buddha, Kiranshastry, Dimitry Miroliubov, Surang and accessed at www.freepik.com.

### Code Accessibility

Task and analysis code, as well as raw data and pre-processed data will be released on publication.

### Experimental Design

Participants performed a sequential decision-making task that required deciding when to select an option on screen, as its reward changed from trial to trial (Figure 1). There were two critical aspects to the task. First, each option had a starting value and a growth rate (the increase in reward per trial). These properties were varied independently, resulting in unique options that reached a 500-point reward maximum after a different number of trials. After reaching the 500-point maximum, the option reward began to decline, encouraging participants to select the option on the single trial when it was at maximum reward. Second, the visual appearance of the option remained the same from trial to trial, despite changes in its reward value.

**Figure 1.**
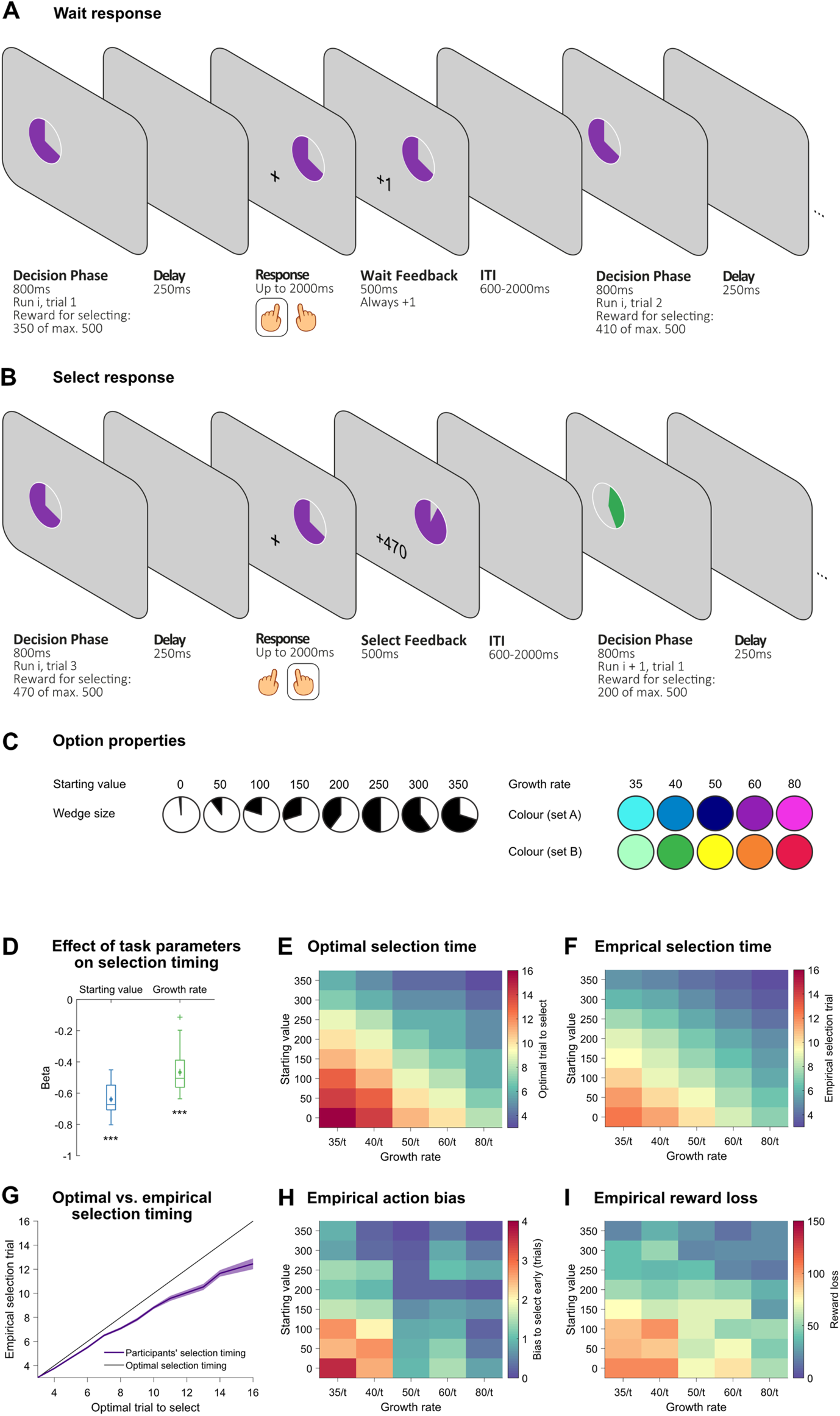
**A-B**: Task design. Participants were presented with an option that had a specific starting value (wedge size) and growth rate (colour). **A:** Participants could choose to let the option’s reward increase, by selecting the blank side of the screen during the response phase. The option reward was then updated by the growth rate and the participant performed another trial with the same item. Importantly, the same starting value and growth rate information were presented for each trial within a run, even though the underlying option reward was changing. **B:** Participants could choose to end the trial run by selecting the side of the screen with the option during the response phase. Participants then received the current option reward and started a new trial run, with a new option. Option rewards increased linearly up to their maximum value (500 points), after which value decayed exponentially if participants continued to make wait responses. Trial runs lasted up to a maximum of 16 trials. **C:** Option properties. Options could have one of eight starting values, ranging from 0-350 points.

A single option was presented in each trial run. The option was a circle stimulus with an approximate visual angle of 3.79° (150×150 pixels), calculated based on an approximate viewing distance of 60cm. The option contained a coloured wedge, the size of which indicated its starting value and the colour of which indicated its growth rate. On each trial in the run, participants decided between waiting and selecting the option. The run continued until participants decided to select, up to a maximum of 16 trials. Each trial had three phases: a decision phase, where participants were instructed to decide whether to wait (allowing the option’s latent reward to update) or select the option (to gain the latent reward); a response phase, where participants made a manual response to indicate their choice; and a feedback phase. In the decision phase, the option was presented for 800ms on the left or right of the screen, with the centre of the option being presented at an approximate visual angle 3.16° (125 pixels) above and 3.16° to the left or right of the screen’s centre. The decision phase was followed by a 250ms blank delay. The response phase began with the presentation of a central fixation cross, with the option appearing randomly on the left or right of the screen.

Participants were instructed to press the button on the side where the option was presented to select it (e.g. left button if the option was presented on the left) and to press the button on the opposite side to wait (e.g. left button if the option was presented on the right). Since the option appeared randomly on the left or right of the screen, independent of where it had been presented in the decision phase, participants were prevented from preparing a motor response prior to the response phase, allowing us to decouple decision and response information in the neural analyses. In the feedback phase, participants received 1 point for a wait response and the option’s latent reward for a select response (0-500 points). Feedback presentation lasted 500ms. Each trial was followed by an inter-trial interval (ITI). Durations for each ITI in a run were taken from a shuffled ITI vector. The vector contained 8 durations from 500-1200ms, which were spaced 100ms apart. The vector contained two repetitions of each duration.

The reward associated with each option could grow to a maximum of 500 points. Participants were told about the maximum, and asked to select each option when the maximum was reached. The reward was computed with the following linear function: *r* = *s* + *g* (*t* − 1), where *r* is the reward, *s* is the starting value, *g* is the growth rate and *t* is the trial number within the run. Eight different starting values were used in the study (0, 50, 100, 150, 200, 250, 300, 350). On trial runs where the starting value was 0, a coloured wedge corresponding to 7.5 points was used so that participants could see the coloured wedge and determine the growth rate. Five different growth rates were used in the study (35, 40, 50, 60, 80). These were selected to maximize behavioural variability, producing an even spacing in optimal trial on which to selection the option (16, 14, 12, 10, 8 trial steps), when the option started from a value of 0 points. For each participant, two colours were used per growth rate so that abstract growth rate information, independent of option colour, could be decoded from neural recordings. The mapping between specific colours and specific growth rates was pseudorandomised across participants, who were assigned to one of four stimulus groups. All groups had two colours per growth rate, but specific colours corresponded to different growth rates for each group. On trials where the updated option value exceeded 500 points, the option value was rounded down to the 500-point maximum. The largest amount that the 500 points would have been exceeded by a condition without rounding down was 70 points. If participants did not select the option on the trial where it reached its maximum, the reward associated with the option would begin an exponential decay given by the equation: *r* = m(*e^−λt^*), where *r* is the reward, *m* is the maximum value of 500 points, *e* is Euler’s number, *λ* is a decay constant set at 0.5 and *t* is the number of trials elapsed since the option reached the 500-point maximum. This equation meant that each option had just one trial at maximum reward in a run. It was designed to be both distinct from the linear growth function above and the uniform across all conditions, so that participants could not gain information about an option through over-waiting and there would be a strong incentive for participants to end the run once the maximum was reached. To motivate participants to earn as many points as possible, every five percent of the maximum task points earned resulted in a bonus item being unlocked (such as a rocket ship, balloons or an electric guitar), each of which corresponded to additional £1 bonus payment.

Participants completed a training sequence of 40 trial runs prior to the main experiment. If a participant scored below 70% or wished to do more practice, they completed an additional 40-80 trial runs. Eight participants completed 40 practice runs, twenty-one participants completed 80 practice runs and one participant completed 120 practice runs. The main purpose of the practice was to learn the correspondence between the 10 colours and the five growth rates. To facilitate this, participants were presented with an on-screen colour legend during task practice, which indicated the colours for each growth rate. The colour legend was not presented during the main task. Before beginning the main experiment, participants performed a colour ranking task, in which they had to rank the colours on the basis of their growth rates (from 1=lowest growth rate to 5=highest growth rate). If any ranking was incorrect, the task practice was repeated; participants needed to score 100% on the ranking task before starting the main experiment. During EEG recording, participants performed 240 trial runs, with equal numbers of runs for each starting value/growth-rate combination. Equal number of runs were also performed using the two colours for each growth rate. As participants decided when to select the option during each trial run, different numbers of individual trials were recorded per participant. On average, we recorded 1660 trials for each participant (min=1412, max=1898).

### EEG Pre-processing

EEG data were down-sampled from 1000 to 250Hz and filtered using 40Hz low-pass and 0.01Hz high-pass filters. For each participant, channels with excessive noise were identified by visual inspection and replaced via interpolation, using a weighted average of the surrounding electrodes. Data were then re-referenced by subtracting the mean activation across all electrodes from each individual electrode at each time point. Data were divided into epochs from −1 to +5 seconds from the first option onset of each trial. Epochs containing artefacts (such as muscle activity) were rejected based on visual inspection. Data were then subjected to an Independent Component Analysis. Structured noise components, such as eye blinks, were removed, resulting in the data set used for subsequent analyses.

### Behavioural Analyses

To test whether selection timing was sensitive to task variables, we used linear regression. We first tested whether starting value and growth rate could be used to predict the trial on which the option was selected within the run. This used the following equation to estimate one beta per predictor for each participant: 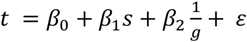, in which *t* is the trial where the option is selected, *β*_0_ is a constant, *s* is the starting value, *g* is the growth rate and *ε* is the residual error. The 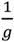 in this expression arises from the fact that the reward, *r* = *s* + *g*(*t* − 1). This in turn means that the trial on which an option reaches maximum reward, **r*_max_*, can be calculated as: 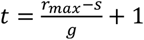. While appropriate due to the reward dynamics in this task, 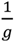 has the effect of inverting growth rate values in the regression. This means that higher growth rates are represented with lower numbers. We therefore multiplied the subsequent *β*_4_ estimates by −1, to correctly interpret the direction of the effect. Once one beta had been estimated per predictor for each participant, the 32 participant betas for starting value and the 32 participant betas for growth rate were tested against zero in separate two-tailed t-tests. In further exploratory analyses, we found that participants tended to select options earlier than would be optimal for the slowest trial runs (i.e. those that took the most trials to reach maximum reward). We use the term action bias to refer to this tendency to select earlier than would be optimal for reward maximisation. The action bias was computed by subtracting the optimal trial to select the option within the run from the empirical trial on which the option was selected. The vector was then multiplied by −1 to convert the action bias to a positive scale. The following regression equation was used for the action bias analysis: *b* = *β*_2_ + *β*_3_*t* + *ε*, where *b* is the action bias, *β*_2_ is a constant, *t* is the optimal to trial on which to select the option to gain maximum points and *ε* is the residual error. The resulting 32 participant *β*_3_ estimates were tested against zero using a two-tailed t-test. Using a similar approach, we explored whether the amount of reward lost on each trial run could be predicted from the optimal number of trials to wait for each condition. The reward loss was computed by subtracting the amount of reward earned on each run from the total reward possible on that run. The total reward possible was the 500-point maximum plus one point for each wait decision needed to reach the optimal selection time. The reward loss vector was then multiplied by −1 to convert it to a positive scale. The following regression equation was used for the reward loss analysis: *l* = *β*_2_ + *β*_3_*t* + *ε*, where *l* is the reward loss, *β*_2_ is a constant, *t* is the optimal to trial on which to select the option to gain maximum points and *ε* is the residual error. The resulting 32 participant *β*_3_ estimates were tested against zero using a two tailed t-test. Trial runs in which a select response was not made within 16 trials were excluded from all behavioural analyses. Dependent and independent variables were z-scored prior to running all regressions. Exploratory analyses were corrected for multiple comparisons using the Bonferroni-Holm correction (Holm, 1979). The correction was applied within each analysis. For the one exploratory comparison between starting value and growth rate regressors, the alpha threshold was unchanged. For the action bias and reward loss exploratory analyses (which both used the optimal trials to wait as a predictor), the alpha threshold was corrected for two exploratory tests.

### EEG Analyses

We conducted a series of multivariate EEG analyses to understand the neural processes underpinning task performance. The analyses aimed to identify EEG signals that tracked three sets of task variables: (1) externally presented variables (starting value and growth rate), (2) internally represented variables (distance-to-goal and latent reward) and (3) action variables (distance-to-select, latent reward to select and select versus wait response). We predicted that participants would track an option’s latent reward and use this information to decide when to select the option, a strategy that would be reflected as strong latent reward encoding.

### Cross-validated neural decoding

To test the encoding of each task variable, we used a cross-validated decoding approach. We will describe the approach in general terms first and provide details about the specific analyses in subsections below. To decode a specific task variable, we selected data from the relevant trial (e.g. trial 1 within the run). Trials that involved a select response or no response were excluded. Data were baselined from 250ms to 50ms before trial onset and channel demeaned prior to all analyses. The analysis then ran through a series of train and test folds. In each fold, a maximum of one trial per condition was held out as test data. As an example, the test trials for one starting value fold could contain a total of 8 trials with one trial for each starting value. The remaining trials were used as the training data for that fold. The number of trials for each condition was balanced in the training fold by selecting a random subsample of non-test trials for each condition. The number of trials in each random subsample was one trial less than the lowest number of trials across the conditions, for trial positions 1-3. Analyses primarily focused on trial positions 1-3 because these were trials in which the majority of conditions (39/40) had not yet resulted in a decision to select, under the optimal selection timing. Balancing the trial numbers across conditions and trial positions meant the decoder would not be biased towards a particular condition and that decoding results could be compared between trial positions.

Once the training data and test data were organised for a fold, the training trials were averaged for each condition. This produced a channels × time matrix that reflected the mean scalp topography for each condition. Averaging over training trials within each condition has been shown to improve decoding accuracy (Grootswagers et al., 2017; Isik et al., 2014). In EEG analyses based on voltages, it has the additional benefit of averaging out non-phase-locked background oscillatory activity. Following this step, each held out test trial was compared to the average topographies. To make this comparison, Mahalanobis distances (MDs) were calculated at each time point between the test trial (a vector of 61 channel values) and each condition mean (each a vector of 61 channel values). We selected the MD to measure decoding because it explicitly accounts for covariance structure in the data. This makes it well suited to EEG data, where channel values tend to be highly correlated. The MD between the test vector (pattern A) and a condition vector (pattern B) was computed as: 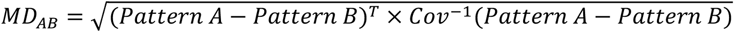, where Pattern A-Pattern B is the difference between topographies, T is the transpose and Cov^-1^ is the inverse of the channel covariance matrix. The channel covariance matrix was estimated using within-condition error, meaning that trials were condition demeaned prior to estimating the covariance (Walther et al., 2016). The channel covariance estimate also included a shrinkage estimator (Ledoit & Wolf, 2004), which downweights noisy covariance estimates.

Once MDs had been computed between each test trial in a fold and the condition averages, the MDs for each trial were then entered into a regression. The MDs were used as the dependent variable and a vector of predicted condition distances were used as the independent variable. The predicted distance vector assumed a linear increase in MD for more dissimilar conditions. For example, a test trial with a starting value of 0 would have a distance vector of [0,1,2,3,4,5,6,7], indicating there is no expected difference between the test trial and the condition mean for a starting value of 0, but a maximal difference between the test trial and the condition mean for a starting value of 350. Both dependent and independent variables were z-scored prior to running the regression, which was performed at each time point. Once this regression procedure had been completed for each test trial in the fold, the next analysis fold began, using a new set of test trials. This process continued until all trials had been used as a test trial once, resulting in a trials × time matrix of regression coefficients (betas). To ensure the beta estimates were robust, the entire analysis procedure was repeated 50 times, with eligible data being randomly assigned to different folds on different repetitions. The betas were then averaged over these repetitions and over trials, providing a time course of beta that we interpret as neural coding of a given task variable. Once this process was completed for trial position 1 data, it was applied to data from trial positions 2 and 3.

To assess significant decoding of a task variable, beta estimates for each trial position were averaged over the decision phase, from trial onset (0s) to the offset of the option stimulus (800ms) and tested against 0 using two-tailed t-tests, with Bonferroni-Holm (BH) adjusted alpha thresholds to correct for the three trial positions (Holm, 1979). To test differences in decoding between trial positions, the beta averages were compared for each pair of trial positions (i.e. 1 vs 2, 1 vs 3, 2 vs 3) using two-tailed t-tests, with adjusted alpha thresholds to correct for three between-position tests (Holm, 1979). When examining the decoding time course for a task variable, trial numbers were matched across conditions during the analysis but not across trial positions to increase power. Time courses were smoothed with a 20ms Gaussian kernel prior to non-parametric cluster-based permutation testing, which was used to correct for multiple comparisons (Maris & Oostenveld, 2007; Hall-McMaster et al., 2019; Sassenhagen & Draschkow, 2019).

### Presented variables

#### Starting value

When decoding information related to the starting value of the choice item (i.e. the arc length), there were 8 possible starting value conditions (0, 50, 100, 150, 200, 250, 300 and 350). For time-averaged analyses, where trial numbers were balanced across the conditions and trial positions, the average number of trials per condition across participants was 19.5 with a standard deviation of 2.94. When examining the time course at trial position 1, where starting value coding was strongest, matching trial numbers at trial position resulted in a mean of 25.7 trials per condition and a standard deviation of 2.46.

#### Growth rate

When decoding abstract growth rate information, the analysis was adjusted to leverage the fact that there were two colours per growth rate. This allowed us to decode abstract growth rate information that could not be attributed to the colour of the option. Trials using the first colour set were assigned to data split A and trials using the second colour set were assigned to data split B. We then maintained a strict separation between data splits A and B in our decoding analyses. Data split A was only used as training data and data split B was only used as test data (and vice versa). There were 5 possible growth rate conditions (35, 40, 50, 60, 80). The regression for each test trial therefore used a vector of predicted condition differences that had 5 values as the independent variable. For time-averaged analyses, where trial numbers were balanced across conditions, trial positions and colour set, the average number of trials per condition across participants was 17.4 with a standard deviation of 1.29. When examining the time course at trial position 1, where growth rate coding was strongest, matching trial numbers for each growth rate resulted in a mean of 19.9 trials per condition and a standard deviation of 2.05.

### Latent variables

#### Latent reward

For latent reward decoding, data were separated into 50-point bins, starting from 0 points (bin 1=0-49 points, bin 2=50-99 points, bin 3=100-149 points, bin 4=150-199 points, bin 5=200-249 points, bin 6=250-299 points, bin 7=300-349 points, bin 8=350-399 points, bin 9=400-449, bin 10=450-500 points). To make the analysis comparable with our analysis of distance-to-goal (described next), we used a sliding range of 6 latent reward bins (bins 4-9 at trial position 2 and 5-10 at trial position 3) that corresponded approximately to the goal distances used in distance-to-goal decoding. Note that at trial position 1, the reward value of the choice item is the starting value. The item’s latent reward value only differs from starting value after trial 1, which is why this analysis only considers trials 2 and 3. Statistical tests were accordingly corrected for two comparisons (Holm, 1979). For time-averaged analyses, where trial numbers were balanced across bins and trial positions, the average trials per condition was 21.4, with a standard deviation of 1.77. The vector of predicted conditions differences used in the regression was based on the difference in reward bin number. When examining the time course at trial position two, the average number of trials per condition was 21.9, with a standard deviation of 1.73.

#### Distance-to-goal

This task variable refers to how many trials away the participant is from the optimal trial to select the option. When decoding distance-to-goal information, trials were therefore sorted by the difference between the current trial and the optimal trial to select the option. The analysis used a sliding range of distance-to-goal conditions at each trial position (trial position 1 = goal distances 3-8, trial position 2 = goal distances 2-7, trial position 3 = goal distances 1-6). The sliding range aimed to maintain the same future selection trials in the analysis at each trial position. At trial position 1, the optimal trials to select spanned from trial 4 to trial 9 within the run. Sliding the goal distances meant this was also true at trial positions 2 and 3. This ensured that differences in decoding between trial positions were not due to differences in starting value, growth rate or the optimal trial to select, because these factors were consistent across positions. The choice of which goal distances to use aimed to balance a trade-off between data quality, in which behaviour was increasingly suboptimal further into a trial run (Figure 1G), and power for the regressions between each test trial’s Mahalanobis distances and the vector of expected condition distances. Using more goal distances would increase the number of points for these regressions but it would also include more trial runs in which selection behaviour was increasingly suboptimal. While we aimed to balance this trade-off, the choice of goal distances was still somewhat arbitrary. This made it important to replicate the distance-to-goal coding results in the more carefully controlled RSA analyses (described below). When balancing all trial numbers across the conditions and trial positions, the average number of trials per condition across participants was 11.3488 with a standard deviation of 3.1172. When examining the time course at trial position 2, where distance-to-goal coding was strongest, matching trial numbers across distance-to-goal conditions resulted in a mean of 16.7188 trials per condition and a standard deviation of 1.0234.

#### Action variables

The previous analyses were performed separately at each trial position. When decoding action variables, trials were pooled across trial positions 3-6 and entered into a single analysis to boost power. The increased trial numbers allowed us to do 10-fold cross-validation, as opposed to the more cumbersome process of holding out 1 trial per condition for each test set in the analyses above. The trials in non-test folds were used as training data. The number of trials from each condition and each trial position were balanced in the training set by taking random subsamples from the non-test folds. To avoid potential reaction time (RT) differences between conditions affecting the decoding results, RTs for each condition in the training set were statistically compared using paired two-tail t-tests. The alpha threshold was adjusted to correct for multiple comparisons using the Bonferroni-Holm correction (Holm, 1979). If two conditions had significantly different RTs, a new random subsample was selected from the available training trials until no significant differences between condition RTs were detected. Only then was the analysis permitted to proceed with calculating multivariate distances between training and test trials. As with previous analyses, we repeated the analysis 50 times and averaged the outputs to ensure stability of the decoding results.

#### Distance-to-select

The distance-to-select refers to the number of trials between the current trial and the trial where a select response is made. For this analysis, select distances of 1-3 were used as conditions. To control for the possibility that the trial immediately before the decision to select was distinct from all other trials prior to the decision to select, we ran a control analysis that only included select distances of 2 and 3. To control for the possibility that our decoding results were driven by different numbers of left and right handed responses, we ran a second control analysis for select distances 1-3, in which we additionally balanced the number of left/right button presses.

#### Reward to select

The reward to select refers to the difference in latent reward on the current trial and the latent reward when the select response is made. We calculated this difference for each trial in each run. The reward to select values were then separated into 50-point bins (bin 1=0-49 points, bin 2=50-99 points, bin 3=100-149 points). Reward to select bins of 1-3 were used as conditions in the analysis.

#### Wait vs select

This was same as the distance-to-select procedure, decoding select trials vs the previous wait trial (i.e. distance-to-select 0 vs. distance-to-select 1).

### Representational Similarity Analysis

We used RSA (Kriegeskorte et al., 2008) to control for the possible influence of extraneous task variables when decoding distance-to-goal and latent reward. The logic of this approach was to regress out neural coding of starting value, growth rate and latent reward information, before testing for distance-to-goal coding. In a similar manner, we wished to regress out neural coding starting value, growth rate and distance-to-goal, before testing for latent reward coding. If decoding is still significant for the variable of interest after removing the influence of other task variables, this confirms that our earlier cross-validated decoding result is not being driven by correlations between task variables. If decoding for a variable of interest is no longer significant, it indicates that our earlier cross-validated decoding result could have been driven by another task variable.

We first focused on the analysis of data from trial position 1. We iterated through each starting value-growth rate-colour set combination, yielding 80 conditions in total (8 starting values × 5 growth rates × 2 colour sets). The 80 conditions were repeated 3 times each in the experiment. However, a condition could have less than 3 trials available for analysis at a given trial position due to trials being rejected during pre-processing or due to trials containing a select response. To ensure balanced trial numbers in the RSA, we included conditions with all 3 available trials at a given trial position. This resulted in 15/80 conditions being excluded on average at trial position 2, where latent reward and distance-to-goal were detected in earlier decoding analyses. No significant differences were found between the excluded and included conditions on trial 2, based on mean starting value (*t*(31)=−0.508, *p*=0.615) or mean growth rate (*t*(31)=−1.015, *p*=0.318). For each eligible condition, we averaged over trials to get the mean scalp topography at each time point. MDs between each pair of conditions were computed at each time point, resulting in a participant-specific representational dissimilarity matrix (RDM) for each time point. Next, we constructed a set of model dissimilarity matrices to reflect the expected dissimilarity structure based on different task variables. Model matrices were calculated by computing the difference between each condition pair of a given task variable (starting value, growth rate, distance-to-goal, latent reward, colour set used to indicate the growth rate). Note that unlike the decoding analyses, here the latent reward model was not based on the difference in reward bins but the exact difference in latent reward between conditions.

The first stage regression aimed to remove information that was extraneous to the variable of interest from the data RDMs at each time point. When examining latent reward as a variable of interest, model RDMs for the extraneous variables (starting value, growth rate distance-to-goal, number of trials used to compute the condition average, growth rate colour set), and the data RDM were transformed into vectors and z-scored. For each time point, the data RDM was used as the dependent variable and model RDMs for the extraneous variables above were used as independent variables in a multiple regression, which included a constant regressor. In a second stage regression, the residual variance from the first stage was used as the dependent variable latent reward RDM (transformed into a vector and z-scored) was used as the independent variable. This resulted in a time course of regression coefficients (betas) for latent reward coding that did not reflect a linear influence of the extraneous variables above. The analysis procedure was repeated for trial positions 2 and 3. The procedure was the same when distance-to-goal was the variable of interest, except that the distance-to-goal model RDM was used as the independent variable during the second stage regression and the latent reward model RDM was included as an extraneous variable in the first stage regression. Like the cross-validated decoding analysis, latent reward coding was only tested for trial positions 2 and 3 because latent reward and starting value were indistinguishable for trial position 1. Statistical assessment of the decoding strength was tested in the same way as cross-validated decoding analyses. Beta coefficients for the variable of interest were averaged over the decision phase (0-800ms).

#### Spatial RSA

Having identified distance-to-goal encoding at trial position 2, we performed an exploratory analysis to test whether this information was encoded in a specific subset of electrodes. To do so, we repeated the distance-to-goal coding RSA analysis three times, once using a subset of 18 frontal electrodes (F7, F5, F3, F1, Fz, F2, F4, F6, F8, FT7, FC5, FC3, FC1, FCz, FC2, FC4, FC6, FT8), once using a subset of 18 central electrodes (T7, C5, C3, C1, Cz, C2, C4, C6, T8, TP7, CP5, CP3, CP1, CPz, CP2, CP4, CP6, TP8) and once using a subset of 17 posterior electrodes (P7, P5, P3, P1, Pz, P2, P4, P6, P8, PO7, PO3, POz, PO4, PO8, O1, Oz, O2).

### Relationships between task performance and neural coding

#### Models of selection timing

To provide broad insights about the cognitive strategy used to guide selection behaviour, we performed an exploratory cross-validated model fitting analysis. The aim of this analysis was to arbitrate between high-level explanations for participants’ bias to select early on long trial runs. We reasoned that early selection could arise for two broad reasons. First, participants could assign subjective cost to the number of steps waited, which would reduce the maximum reward that could be gained on a trial run. This lower subjective maximum would be reached sooner than the true maximum and thereby result in earlier selection. Second, participants could have a biased representation of the starting value or growth rate, either of which would result in an inaccurate estimate of the latent reward on each trial. Such a bias would have a compounding effect on the latent reward estimate, resulting in more inaccurate reward estimates as more trials are waited. This in turn would result in an increasing bias to select early as the optimal selection time increased.

To test these broad accounts, we created a series of different models. Each model was based on the fact that the task’s reward dynamics followed the equation: *r* = *s* + *g*(*t* − 1), where *r* is the reward, *g* is the growth rate and *t* is the current trial within the run. This meant that the optimal trial on which to select the option, *t*\*, could be expressed as: 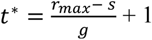, where **r*_max_* is 500 points. To create the cost of waiting model, we added a hyperbolic delay discounting factor to the maximum reward (see van den Bos & McClure, 2013) resulting 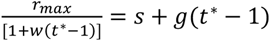, where *w* is a free parameter for the cost of waiting. This equation can in turn be rearranged into the form: *gwn*^2^ + (*g* + *sw*)*n* + (*s* − *r_max_*) = 0, where *n* = *t*\* − 1. The equation can then be solved for n using the quadratic formula: 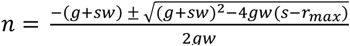, after which *t*\* is calculated as *t*\**=* n + 1. The biased starting value model was formulated as: 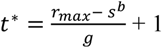, where b is a free parameter for the starting value bias. The biased growth rate bias model was formulated as: 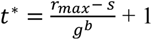, where b is a free parameter for the growth rate bias. An exponential bias was used in these models, as opposed to a multiplicative or additive bias, to account for the non-linear bias observed in empirical selection times (Figure 1G). We also created models with each pairwise combination of the free parameters (wait cost, starting bias, growth bias) and one model with all three free parameters.

The performance of each model was evaluated in a cross-validation procedure. For each participant, data were randomly divided into 10 folds. On each iteration of the analysis, one fold was held out as test data and the remaining folds were used as training data. The best fitting values of the free parameter/s in a model were selected by minimising the sum of the squared error. The minimum and maximum possible parameter values for the waiting cost parameter were constrained to 0 and +30. For the starting value and growth rate bias parameters, these were 0 and +5. The best fitting parameter values were then used in each model equation, to predict the trials waited until a selection response was made in each of the held out test runs. The model predictions were rounded up to the nearest integer to account for trials being discrete units. The model was then scored for that test fold by calculating the mean squared error (MSE) between the predicted trials waited and the actual trials waited.

This process was repeated until each model had a MSE score for each test fold. The procedure was repeated 50 times with random subsamples of trials in each fold to ensure stable results. Within each subsample, the different models were trained and tested on the same trials to provide a fair comparison of performance. Scores were then averaged across the test folds and the subsamples. For each participant, we next calculated the difference in MSE between each model and the cost of waiting model. Once we had an MSE difference score for each model for each participant, we calculated bootstrapped 95% confidence intervals for the mean difference in MSE. The differences in MSE were assessed using Bayesian paired t-tests, to quantify the strength of evidence that the wait cost model had lower error than the other models. When examining the predictions of the three best models against the optimal selection model (Figure 4A-C), we added 1 to both the predicted number of trials waited and the actual number of trials waited, thereby examining the model and participant data in terms of the trial on which the option was selected.

#### Relationship between neural coding and model parameters

The model parameters for the cost of waiting model were averaged over the test folds. These values were then entered into a series of Spearman correlations with the decoding strength of different task variables, taken from the cross-validated decoding analyses described above. These task variables included the decoding strength, averaged over the decision phase (0-800ms), for the starting value on trial 1, growth rate on trial 1, distance-to-goal on trial 2 and latent reward on trial 2. The task variables also included the decoding strength, averaged over the response phase (1050-2000ms), for the distance-to-select and wait versus select decoding. The alpha threshold was adjusted for 6 exploratory tests using the Bonferroni-Holm correction (Holm, 1979). Correlations between these model parameters and the neural data could potentially provide converging evidence for a cost of waiting account. Specifically, we reasoned that if the cost of waiting parameters reflected a subjective waiting cost, they should correlate with a decision variable needed to evaluate when a select response should be made, such as distance-to-goal information. To validate the correlation results in a model-free manner, we repeated the correlations but substituted the cost of waiting parameter for each participant with their average action bias (the average number of trials earlier participants selected the option than the optimal linear model).

## Results

To investigate whether decisions about when to select an option arise from latent reward or distance-to-goal tracking, 32 participants completed a wait-select task (Figure 1). On each trial run, participants were presented with one of 40 specific options, defined by a combination of one of eight possible starting values and one of five growth rates. These properties were signalled using a triangular wedge shown on screen, the size of which indicated the starting value and the colour of which indicated the growth rate. On each individual trial within a run, participants decided between waiting and selecting the option. When deciding to wait, the option’s reward increased based on its growth rate, towards a maximum of 500 points. Participants then moved to the next trial, making another wait-select decision. The unique starting value and growth rate combinations meant that different options reached the 500-point maximum after a different number of wait decisions. Once the reward maximum had been reached, continued wait decisions produced an exponential decline in the option’s reward value, forcing participants to carefully time each select decision to gain the maximum number of points. The timing for optimal selection could range from 3 to 16 trials into the run (mean=8, SD=3) and the run could last for a maximum of 16 trials. Critically, the visual appearance of the option on the screen remained the same throughout the trial run, despite changes in its underlying reward value following each wait decision. This meant that participants could not use sensory information to guide decision-making after the first trial, and instead needed to use latent information that had been computed internally, such as the option’s latent reward prospect. When deciding to select the option, participants earned its latent reward and the trial run ended.

Participants had experience with options prior to starting the experiment, to learn the correspondence between growth rates and wedge colours. Participants were also told about the 500-point reward maximum, and asked to select an option when the maximum was reached. Each participant completed 240 trial runs in the main task, averaging 1,660 individual wait-select decisions (SD=126). 61-channel electroencephalograms recorded during decisions were used to conduct a series of cross-validated decoding and multi-stage representational similarity analyses (RSA), to assess latent reward and distance-to-goal tracking, in the lead-up to option selection. The orthogonal manipulation of starting value and growth rate in the task made it possible to dissociate an option’s latent reward from potentially confounding variables in the neural analyses, such as how many trials had passed since the beginning of a run and how many trials remained until an option reached its maximum.

### Decisions to select were influenced by starting value and growth rate

To assess participant choices, we ran a regression that tested whether the number of trials waited before selecting the option was influenced by starting value and growth rate (Figure 1D). The analysis revealed that select decisions showed significant sensitivity to both starting value (mean beta=−0.640, SD=0.101, *p*=8.164-27) and growth rate information (mean beta=−0.467, SD=0.127, *p*=8.655e-20). The direction of these relationships indicated higher starting values and growth rates predicted earlier decisions to select. Participants’ selection timing and optimal selection timing for each starting value-growth rate condition is visualised in Figures 1E-F. To summarise, our behavioural results indicate participant choices were influenced by the reward structure of the task.

### Decisions were biased towards early selection when options were slow to reach the maximum value

Participants showed a bias to select options earlier than would be optimal for reward maximisation, especially in conditions which took longest to reach maximum value (Figures 1G and 1H). To explore this effect, we ran a regression that examined whether the bias to select early grew as optimal selection times increased. This confirmed that participants tended to show a larger bias to select early when an option required more trials to reach the maximum value of 500 points (mean beta=0.440, SD=0.193, Bonferroni-Holm (BH) corrected *p*=2.753e-13, correction applied for the six exploratory tests in this section). The bias to select early on slow trial runs resulted in a loss in points compared to the optimal selection timing. The reward loss could be predicted from the optimal selection times (mean beta=0.271, SD=0.131, BH corrected *p*=2.284e-12), indicating larger reward losses for items that were slower to reach the maximum value. The optimal selection times, used as the independent measure in these regressions, were a function of an option’s starting value and growth rate. We therefore ran follow-up regressions, testing whether starting value and growth rate both influenced the suboptimality measures. This was the case for the action bias (starting value beta=−0.288, SD=0.151, BH corrected *p*=1.523e-11; growth rate beta=−0.274, SD=0.175, BH corrected *p*=1.024e-9) and the reward loss (starting value beta=−0.265, SD=0.099 BH corrected *p*=4e-15; growth rate beta=−0.115, SD=0.109, BH corrected *p*=1.520e-6), indicating decisions were more suboptimal for options with lower starting values and growth rates.

Starting value was indicated by the size of a wedge shown on screen (with wedge size oriented randomly on each trial run). The option could have one of five growth rates, which determined the increase in an option’s latent reward per wait response. Growth rates included 35, 40, 50, 60 and 80 points per wait trial, which were selected to maximise variability in the optimal select trial from a starting value of 0. Two colours were used per growth rate for each participant. The two colour sets were used interchangeably during the task. One trial run could use a colour from set A and the next could use a colour from set B, or vice versa. **D:** Regression coefficients showing the influence of starting value and growth rate on the timing of option selection (trial on which the option was selected within the trial run). The lower middle and upper horizontal bars within each plot indicate the 25^th^ percentile, the median and the 75^th^ percentile respectively. Coloured circles within each box show the mean across participants and vertical lines extending from these circles show the standard error of mean. Vertical whiskers extending from each box indicate the most extreme upper and lower values within 1.5 times the interquartile range. Values outside this were deemed outliers and are indicated with a + symbol. *** above each condition indicates regression coefficients are significantly different from zero at p<0.001. **E-F:** Summary of selection timing for each starting value-growth rate combination. **E:** The optimal trial on which to select each option to get the maximum number of points. **F:** The median trial on which participants selected each option. **G:** The trial on which participants selected the option (y-axis) as a function of the optimal trial to select the option in order to gain maximum reward (x-axis). The black diagonal provides a reference line for an optimal agent and the purple line shows mean participant data. Shading around the purple line shows the standard error of mean. **H-I:** Summary of performance biases for each starting value-growth rate combination. **H:** The difference between the optimal and empirical selection timing (the action bias). **I:** The difference between the median number of points earned and the maximum number that could be earned (the reward loss).

### Starting value and growth rate were encoded in neural activity patterns

Having established that participant choices were sensitive to the reward structure of the task, we performed a series of decoding analyses to test whether task information was encoded in patterns of neural activity. Unless otherwise specified, decoding was conducted on the first three trials in a run, where almost all starting-value/growth-rate combinations (39/40) had not yet resulted in a decision to select under the optimal timing. The analyses balanced the number of trials from each condition when training the decoder and used repeated subsampling to ensure stability of the results (see Methods). We first found that information related to the starting value of the choice item (the wedge size) could be decoded during the decision phase (0-800ms following trial onset) during the first three trials within the trial run (Figure 2A). Starting value decoding was numerically highest on the first trial within a run (mean beta=0.018, SD=0.020, *t*(*31*)=5.107, BH corrected *p*=4.739e-5) and decreased on the second (mean beta=0.010, SD=0.013, *t*(31)=4.412, BH corrected *p*=2.998e-4) and third trials with the run (beta mean=0.008, SD=0.019, *t*(31)=2.304, BH corrected *p*=0.028). Despite this numerical reduction, we did not detect significant differences in the decoding strength for starting value related information between trial positions 1 and 2 (*t*(31)=1.685, BH corrected *p*=0.240), positions 1 and 3 (*t*(31)=1.809, BH corrected *p*=0.240 or positions 2 and 3 (*t*(31)=0.727, BH corrected *p*=0.472). When visualising the full time course from 0-2000ms during the first trial (Figure 2B), where starting value decoding was numerically strongest, we detected starting value information across most of the trial, from early in the decision phase into the response phase (window tested=0-2000ms, first significant cluster window=148-1108ms, cluster corrected *p*=3e-04; second cluster window=1140-1364ms, cluster corrected *p*=0.042; third cluster window=1680-2000ms, cluster corrected *p*=0.035). It is important to note that because starting value is coupled to the wedge size, these decoding results could reflect a combination of starting value and perceptual factors like shape and contrast, particularly in the early stimulus-evoked period of the trial (∼100ms). It is equally important to note that separating these factors is not critical for the present study, in which the main aim was to decode latent reward information over time, while dissociating latent reward information from other task factors.

**Figure 2.**
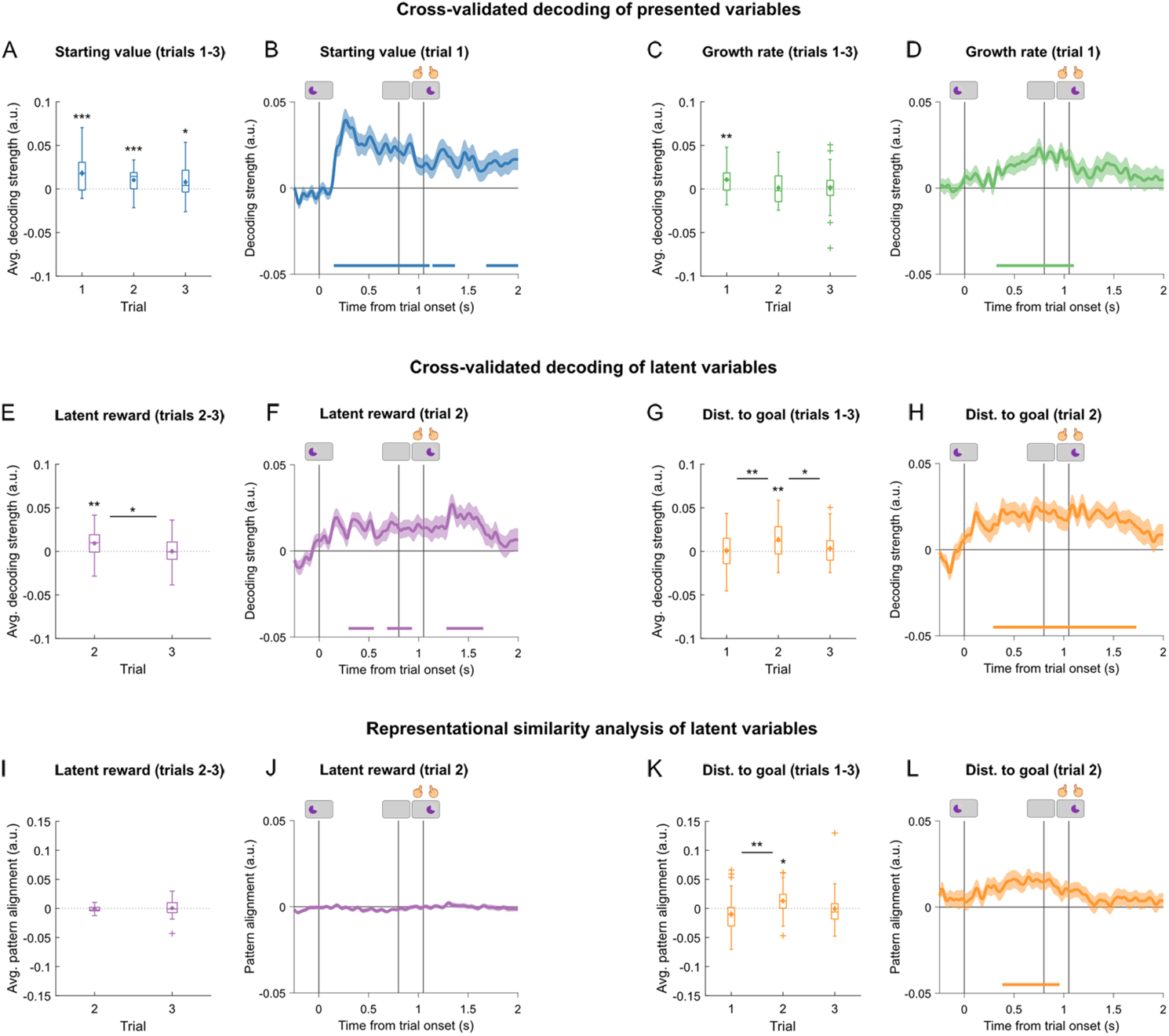
Neural decoding of presented and latent variables. **A:** Average decoding strength for starting value information (wedge size) during the decision phase (0-800ms), on the first three trials within a run. **B:** Time course for starting value decoding for the first trial within a run (0-2000ms from trial onset). **C-D:** Abstract growth rate decoding, independent of stimulus colour due to cross-decoding across colour sets. **C:** Average decoding strength for abstract growth rate information during the decision phase (0-800ms), on the first three trials within a run. **D:** Time course for abstract growth rate decoding on the first trial within a run (0-2000ms from trial onset). **E:** Average decoding strength for latent reward, which is the starting value plus the product of growth rate × trials waited so far in the run. This is shown during the decision phase (0-800ms) on the second and third trials in a run. Latent reward is not shown on trial 1 because it would be equivalent to starting value. **F:** Time course for latent reward decoding on the second trial within a run (0-2000ms). **G:** Average decoding strength for distance-to-goal information (i.e. the number of trials from the current trial to the optimal selection trial) during the decision phase (0-800ms), for the first three trials within a run. **H:** Time course for distance-to-goal decoding on the second trial within a run (0-2000ms from trial onset). **I-L:** Multi-stage representational similarity analysis (RSA) of latent task variables. The cross-validated decoding in figures A-H does not control for the influence of other task variables, which could drive the decoding results. The multi-stage RSA in I-L regresses out neural activity related to other task variables, before decoding the variable of interest. This provides a measure of latent variable decoding, independent from other task variables. The pattern alignment measure refers to how much the neural dissimilarity matrix can be predicted from a model dissimilarity matrix that contains expected condition differences. **I-J:** Pattern alignment for latent reward encoding after starting value (wedge size), growth rate (colour) and distance-to-goal have been regressed out of the neural dissimilarity matrix obtained from the EEG signal. **I:** Average pattern alignment for latent reward encoding during the decision phase (0-800ms) for the second and third trials within a trial run. **J:** Pattern alignment time course for latent reward encoding on the second trial within a run (0-2000ms). **K-L:** Pattern alignment for goal distance encoding when starting value, growth rate and latent reward have been regressed out of the EEG signal. **K:** Average pattern alignment for distance-to-goal during the decision phase (0-800ms), for the first three trials within a run. **L:** Pattern alignment time course for distance-to-goal encoding on the second trial within a run (0-2000ms). **A,C,E,G,I,K:** The lower middle and upper horizontal bars within each box indicate the 25^th^ percentile, the median and the 75^th^ percentile respectively. Coloured circles within each box show the mean across participants and vertical lines extending from these circle show the standard error of mean. Vertical whiskers extending from each box indicate the most extreme upper and lower values within 1.5 times the interquartile range. Values outside this were deemed outliers and are indicated with a + symbol. Asterisk symbols above each condition indicate decoding coefficients are significantly different from zero at ***p<0.001, **p<0.01, *p<0.05. Asterisk symbols above a bar that bridges two conditions indicate a significant difference between conditions at ***p<0.001, **p<0.01, *p<0.05. Asterisk symbols are based on Bonferroni-Holm (BH) corrected p-values. **B,D,F,H,J,L:** Vertical lines show onset of the decision phase, delay and response phase respectively. Shading in the decoding time courses show the standard error of mean. Solid coloured lines under time courses indicate when significant decoding is observed (cluster corrected p-values <0.05).

Unlike starting value, which was coupled to the wedge size, each growth rate corresponded to two wedge colours per participant. This meant we could use cross-decoding to dissociate growth rate encoding from perceptually-evoked responses by training the decoder on one colour set and testing it on the second colour set. Neural encoding of abstract growth rate information, independent of stimulus colour, was present during the decision phase on the first trial within a run (Figure 2C) (mean beta=0.011, SD=0.016, *t*(31)=3.701, BH corrected *p*=0.003). Significant growth rate encoding was not detected on the second (mean beta=0.001, SD=0.019, *t*(31)=0.375, BH corrected *p*>0.99) or third trial within a run second (mean beta=0.001, SD=0.022, *t*(31)=0.284, BH corrected *p*>0.99). We observed a numerical reduction in decoding strength following trial position 1, reflecting a strong trend towards reduced growth rate decoding between trial positions 1 and 2 (*t*(31)=2.484, BH corrected *p*=0.056) and positions 1 and 3 (*t*(31)=2.382, BH corrected *p*=0.056). No significant difference in growth rate decoding was observed between positions 2 and 3 (*t*(31)=0.027, BH corrected *p*=0.979). Visualising the full time course from 0-2000ms during the first trial (Figure 2D), growth rate decoding was evident from midway through the decision phase to the beginning of the response phase (window tested=0-2000ms, cluster window=320-1096ms, cluster corrected *p*=9e-4).

To summarise, information related to an option’s starting value (the wedge size) and abstract information about its growth rate were encoded in patterns of neural activity as participants performed the task. In addition, these task variables were most prominent numerically on the first trial within each trial run.

### Latent reward information was initially decodable from neural activity patterns

So far we have verified that the main task variables were encoded on the first trial of each run. However, either variable on its own is insufficient for participants to solve the task. Starting value and growth rate need to be integrated in order to determine when an option has reached its peak reward. We therefore investigated whether latent decision variables – which were purely internal and could be used to guide selection timing - were encoded in neural activity. One such variable is the option’s latent reward value, which changed from trial to trial, and could be internally tracked towards its maximum using starting value and growth rate information (and elapsed time). On the first trial in the run, the latent reward was equivalent to starting value. We therefore focused on trials two and three, separating trials into 50-point latent reward bins for cross-validated decoding (see *Latent Variables* in Methods). Latent reward information was detected on the second trial within the run (Figure 2E, mean beta=0.009, SD=0.015, *t*(31)=3.551, BH corrected *p*=0.003). Latent reward showed a significant decrease in decoding strength between trials two and three (*t*(31)=2.111, *p*=0.043), with no significant latent reward information detected on trial three (mean beta=6.560e-6, SD=0.016, *t*(31)=0.002, BH corrected *p*=0.998). When examining the full time course during trial 2, where latent reward decoding was strongest, we observed a significant cluster at the start of the decision phase (Figure 2F, window tested=0-2000ms, first cluster window=296-548ms, cluster corrected *p*=0.0232; second cluster window=684-932, cluster corrected *p*=0.031; third cluster window=1284-1648ms, cluster corrected *p*=0.009). To summarise, we found initial evidence that latent reward information could be decoded as early as the second trial.

### Distance-to-goal information was initially decodable from neural activity patterns

Another latent variable that could be used to guide selection behaviour is the number of trials until the maximum number of points is reached, a variable we call the distance-to-goal. Distance-to-goal information could not be detected on the first trial in the run (Figure 2G. mean beta=7.586e-4, SD=0.020, *t*(31)=0.213, BH corrected *p*=0.833). Following trial 1, we observed a significant increase in distance-to-goal encoding (*t*(31)=−3.524, BH corrected *p*=0.004), with significant distance-to-goal information overall on trial 2 (mean beta=0.014, SD=0.020, *t*(31)=3.829, BH corrected *p*=0.002). This distance-to-goal information was encoded transiently, as indicated by a significant decrease between trials two and three within the run (*t*(31)=2.389, BH corrected *p*=0.046), with no detectable distance-to-goal information on trial three (mean beta=0.003, SD=0.019, *t*(31)=0.967, BH corrected p=0.682). No significant difference between trials one and three was detected (*t*(31)=−0.502, BH corrected *p*=0.619). When examining the full time course during the second trial (Figure 2H), where distance-to-goal coding was present, distance-to-goal information was represented during most of the trial, from the decision phase into the response phase (window tested=0-2000ms, cluster window=292-1728ms, cluster corrected *p*=1e-4). To summarise, we found that information about when to select the option for maximum reward could be initially decoded on the second trial within a run, after information about the starting value and growth rate had been encoded on trial 1.

### Latent reward could not be decoded when controlling for other task variables

While latent reward and distance-to-goal information could be decoded in the separate analyses above, these variables were correlated because higher latent rewards meant participants were closer to the optimal trial to select (mean Spearman’s Rho=−0.851 for behavioural data on trial position 2). This means that the decoding analyses above could be sensitive to the same underlying information. To dissociate latent reward and distance-to-goal information, we performed multi-stage RSA. The logic behind this analysis was to first regress out multivariate activity related to extraneous task variables and then test whether information about a variable of interest could still be decoded from the neural signal (see Methods). This approach has the benefit of being able to use exact latent reward values rather than bins, using pairwise differences between condition labels as regressors in a multivariate regression. The correlation between latent reward and distance-to-goal information used in the analysis was reduced in the RSA setup (mean Spearman’s Rho=0.546 on trial 2). The sign flip in this correlation, compared to the raw behavioural correlation above, is due to the use of representational dissimilarity matrices (RDMs) in the analysis. The RDMs capture absolute pairwise differences in condition values. The sign flip occurs because the behavioural values are negatively related, but the pairwise differences in values end up being positively related. This means that two trials with a bigger difference in latent reward also tend to have a bigger difference in their distance-to-goal values. When controlling for the possible influence of starting value, growth rate and distance-to-goal using multi-stage RSA, latent reward information could not be detected on trial 2 (Figure 2I, mean beta=−0.001, SD=0.006, *t*(31)=−1.239, BH corrected *p*=0.449) or trial three (mean beta=4.170e-4, SD=0.014, *t*(31)=0.175, BH corrected *p*=0.862). There was no significant difference in latent reward decoding between trials two and three (*t*(31)=−0.584, *p*=0.563). When examining the full time course on trial 2 (Figure 2J), this control analysis did not detect any significant decoding clusters (window tested=0-2000ms, strongest candidate cluster=660-716ms, cluster corrected *p*=0.2381). This did not change when latent reward was tested in a single-stage RSA, in which starting value, growth rate, distance-to-goal and latent reward were included as predictors in the same RSA step (window tested=0-2000ms, strongest candidate cluster=660-716ms, cluster corrected *p*=0.226). To summarise, we found that when accounting for other task variables, latent reward information could no longer be decoded on trial 2 within the run.

### Distance-to-goal could be decoded when controlling for other task variables

By contrast, when controlling for the possible influence of starting value, growth rate and latent reward, the multi-stage RSA did replicate the cross-validated decoding of distance-to-goal (Figure 2K). As with the decoding analysis, no distance-to-goal information was detected during the decision phase on the first trial within the run (mean beta=−0.010, SD=0.034, *t*(31)=−1.683, BH corrected *p*=0.205). On the second trial, however, distance-to-goal information that was independent from starting value, growth rate and latent reward, could be detected (mean beta=0.013, SD=0.025, *t*(31)=2.919, BH corrected *p*=0.019). This information was transient, with no significant distance-to-goal information detected on trial three (mean beta=−8.393e-4, SD=0.032, *t*(31)=−0.151, BH corrected *p*=0.881). The distance-to-goal encoding seen on trial 2 was significantly higher than on trial 1 (*t*(31)=−3.290, BH corrected *p*=0.008), but no significant differences were detected between trials one and three (*t*(31)=−1.180, BH corrected *p*=0.247) or trials two and three (*t*(31)=1.752, BH corrected *p*=0.179). When examining the full time course on trial 2 (Figure 2L), we detected significant distance-to-goal coding that began in the decision phase and extended into the delay phase (window tested=0-2000ms, cluster window=388-900ms, cluster corrected *p*=0.012). This was also the case when distance-to-goal was tested in a single-stage RSA, in which starting value, growth rate, latent reward and distance-to-goal were included as predictors in the same RSA step (window tested=0-2000ms, cluster window=396-900ms, cluster corrected *p*=0.011). To understand more about the distance-to-goal signal on trial 2, we performed two exploratory analyses. First, we ran the analysis separately for trials with high (200-350) and low (0-150) starting values. This indicated that the effect was primarily (though not exclusively) due to trials that involved low starting values (window tested=0-2000ms, largest candidate cluster=628-780ms, cluster corrected p=0.078), rather than high starting values (window tested=0-2000ms, no candidate clusters). No significant differences were detected between the high and low starting value time courses (window tested=0-2000ms, strongest candidate cluster=256-296ms, cluster corrected *p*=0.335). Second, we examined whether the signal was encoded in a specific subset of electrodes. To do so, we ran the analysis separately using 18 frontal electrodes, 18 central electrodes and 17 posterior electrodes. Distance-to-goal coding was found to be strongest when the analysis was restricted to the posterior electrodes (window tested=0-2000ms, cluster window=200-1148ms, corrected *p*=1e-3; central electrodes: strongest candidate cluster=492-528ms, corrected *p*=0.339; frontal electrodes: no candidate clusters). To summarise, distance-to-goal information, unlike latent reward, was encoded in neural activity even when information about other task variables was removed from the signal.

### Information about empirical selection timing was encoded in patterns of neural activity

So far we have shown that, early within the trial run, participants encoded information about when an option ought to be selected in the future. This distance-to-goal signal was encoded transiently on trial 2 but could not be decoded on the following trial. We speculated that this could arise if the neural response to a given condition starts out as a faithful reflection of the optimal distance to reach the maximum number of points. But since participants’ actual choices were suboptimal (Fig 1G-I), their selection plan might progressively reflect their actual selection timing rather than optimal selection timing, especially on trial runs that involve longer optimal wait times. In this scenario, decoding the actual selection behaviour might still be possible on later trials leading up to the choice. We therefore tested whether participants monitored the number of trials until their actual selection response, a variable we call the distance-to-select. This is distinct from the distance-to-goal because it represents when people made their actual selection response, rather than the optimal response for a given condition, and therefore accounts for suboptimal choice behaviour. To maximise power, these cross-validated decoding analyses pooled data from trial positions 3-6, using balanced trial numbers from each, and tested whether participants were 1, 2 or 3 trials away from making a select response.

Distance-to-select information was encoded across trial positions 3-6 (Figure 3A), with a time course that gradually ramped up from the decision phase to the response phase (window tested=0-2000ms, cluster window=472-2000, cluster corrected *p*<1e-4). Reaction times (RTs) were stable from trials 2-10 before a select response (*F*(8,248)=0.388, *p*=0.926, one-way repeated measures ANOVA), but showed significant slowing on the trial before selecting (mean RT on distance 1 trials=452ms, SD=93 vs mean distance 2=435ms, SD=92, *t*(31)=3.720, BH corrected *p*=0.002; mean distance 1 vs mean distance 3=432ms, SD=92, *t*(31)=3.958, BH corrected *p*=0.001; mean distance 2 vs mean distance 3, *t*(31)=1.477, BH corrected *p*=0.150). However, the neural effect reported here was not due to differences in RTs between the selection distances because the procedure subsampled trials to ensure there were no significant differences in RTs across conditions that could bias the decoder. As would be expected when balancing condition RTs at the participant level, RTs from subsampled trials also showed no significant differences at the group level. Specifically, no significant differences were detected between selection distances of one (mean=487ms, SD=102) and two (mean=486ms, SD=102; *t*(31)=1.021, BH corrected *p*=0.532), one and three (mean=485, SD=104; *t*(31)=1.763, BH corrected *p*=0.263), as well as two and three (*t*(31)=1.133, BH corrected *p*=0.532). The only other neural analysis that could, in principle, have been affected by slower RTs on the trial before a select response was the distance-to-goal RSA analysis (Figure 2L). However, significant distance-to-goal encoding was still seen on trial 2, even when trials immediately preceding a select response were excluded from the analysis (window tested=0-2000ms, cluster window=564-876ms, cluster corrected *p*=0.029). In addition to the RT controls, the distance-to-select effect (Figure 3A) was not due to differences in the number of left and right button presses between conditions. A control analysis that additionally matched the number of left and right responses showed the same results (window tested=0-2000ms, cluster window=476-2000ms, cluster corrected *p*<1e-4). To test whether the effect could be driven by the trial before the select response being distinct from all other trials, we repeated the analysis, but only included trials with a distance-to-select of two or three. This confirmed that distance-to-select information was encoded prior to the trial before option selection (window tested=0-2000ms, first cluster window=212-456, cluster corrected *p*=0.040, second cluster window=1216-1452ms, cluster corrected *p*=0.040, third cluster window=1616-2000ms, cluster corrected *p*=0.012).

**Figure 3.**
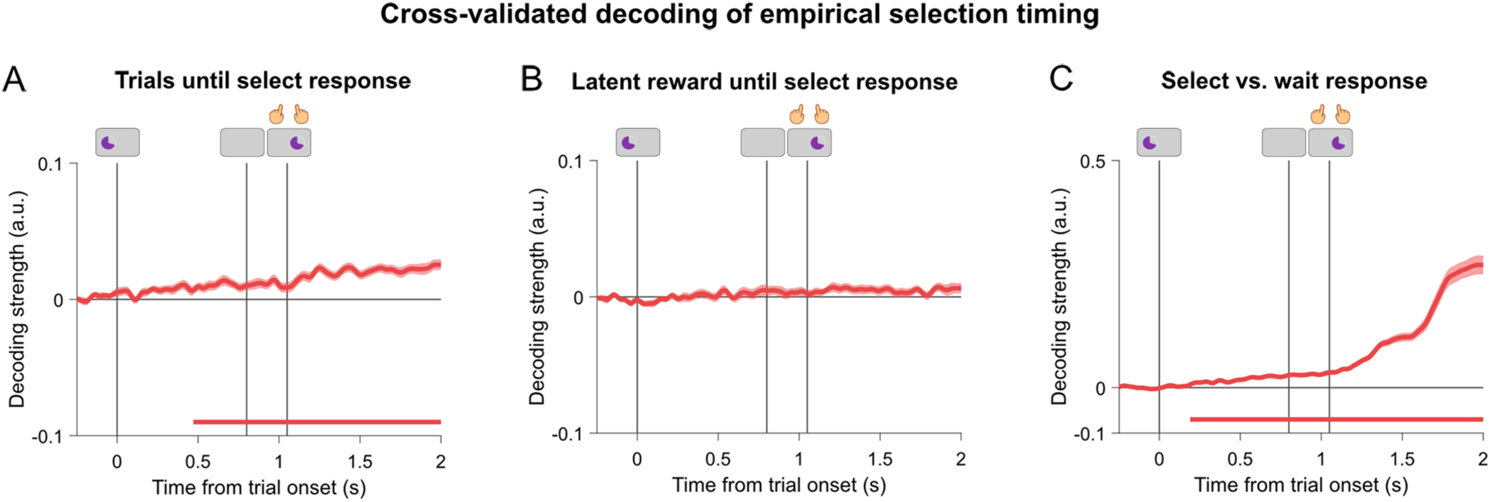
Neural decoding of empirical selection timing. **A:** Decoding time course for the number of wait trials until a select response is made (the distance-to-select). **B:** Decoding time course for the difference between the options current latent reward and its latent reward when selected (the reward to select. **C:** Decoding time course for the decision to select versus the decision to wait on the previous trial. **A,B,C:** Vertical lines show onset of the decision phase, delay and response phase respectively. Shading around the principal line in the decoding time courses show the standard error of mean. Solid coloured lines under time courses indicate when significant decoding is observed (cluster corrected p-values <0.05).

These results suggest that participants performed a mental countdown to time their selection response. However, an alternative explanation could be that participants were counting down the number of points until the reward maximum was reached. This alternative would be consistent with latent reward tracking. To test this possibility, we ran a decoding analysis that examined whether participants were representing the difference in current reward and reward they would get upon selecting the option (the reward to select). The reward to select values were divided into 50-point bins starting from 0 and reward to select bins of 1-3 were used for the analysis (0-150 points away from the select response). We did not detect significant reward to select information during trial positions 3-6 (Figure 3B, window tested=0-2000ms, strongest candidate cluster=44-64ms, cluster corrected *p*=0.597). Having shown participants encoded the trials until their select response, we examined whether decisions to select were neurally distinct from decisions to wait (distance-to-select 0 vs. 1). This revealed a strong select signal that began in the decision phase and became more distinct during the response phase (window tested=0-2000ms, cluster window=192-2000ms, cluster corrected *p*<1e-4).

To summarise, we found evidence indicating that participants tracked the number of trials until their upcoming select response and that this result was not explained through latent reward monitoring.

### Relationship between neural activity and the timing of option selection

So far our neural decoding results point towards a time-oriented task strategy, in which participants evaluated when an option would become most valuable in the future and represented the number of time steps until their upcoming select decision. To understand the relationship between the neural coding results and the timing of option selection, we therefore sought to understand more about the strong bias to select options earlier than would be optimal, when options were slow to reach their maximum value (Figures 1G and 1H). We reasoned that this action bias could emerge for two distinct reasons. One reason would be that the participants discounted an option’s maximum reward based on the number of trials that would need to be waited. This would result in earlier selection because the lower subjective maximum would be reached sooner than the true maximum. A second reason would be that an option’s starting value or growth rate was misrepresented as being higher than its actual value. This would result in earlier selection because the latent reward estimate would become systematically higher than its true value with longer trial runs. When comparing the cross-validated performance between models that could potentially account for the action bias (Figure 4D, see Methods), we found that the model discounting an option’s maximum reward with a waiting cost made more accurate predictions about selection timing than models including a biased representation of the starting value (average difference in mean squared error (MSE)=−2.011; CIs=[−2.687 −1.483], BF_10_=6.131e4), a biased representation of the growth rate (average difference in MSE=−0.164, CIs=[−0.277 −0.079], BF_10_=12.854) or both (average difference in MSE=−0.1341, CIs=[−2.498e5 −2.430e5], BF_10_=5.740). We also found the cost of waiting model did not reliably make more accurate predictions when additional free parameters were added to it, including a bias in the starting value (average difference in MSE=0.015, CIs=[−0.039 0.068], BF_10_=0.215), a bias in the growth rate (average difference in MSE=−0.007, CIs=[−0.047 0.038], BF_10_=0.197) or both (average difference in MSE=0.030, CIs=[−269.353 −259.400], BF_10_=0.242).

**Figure 4.**
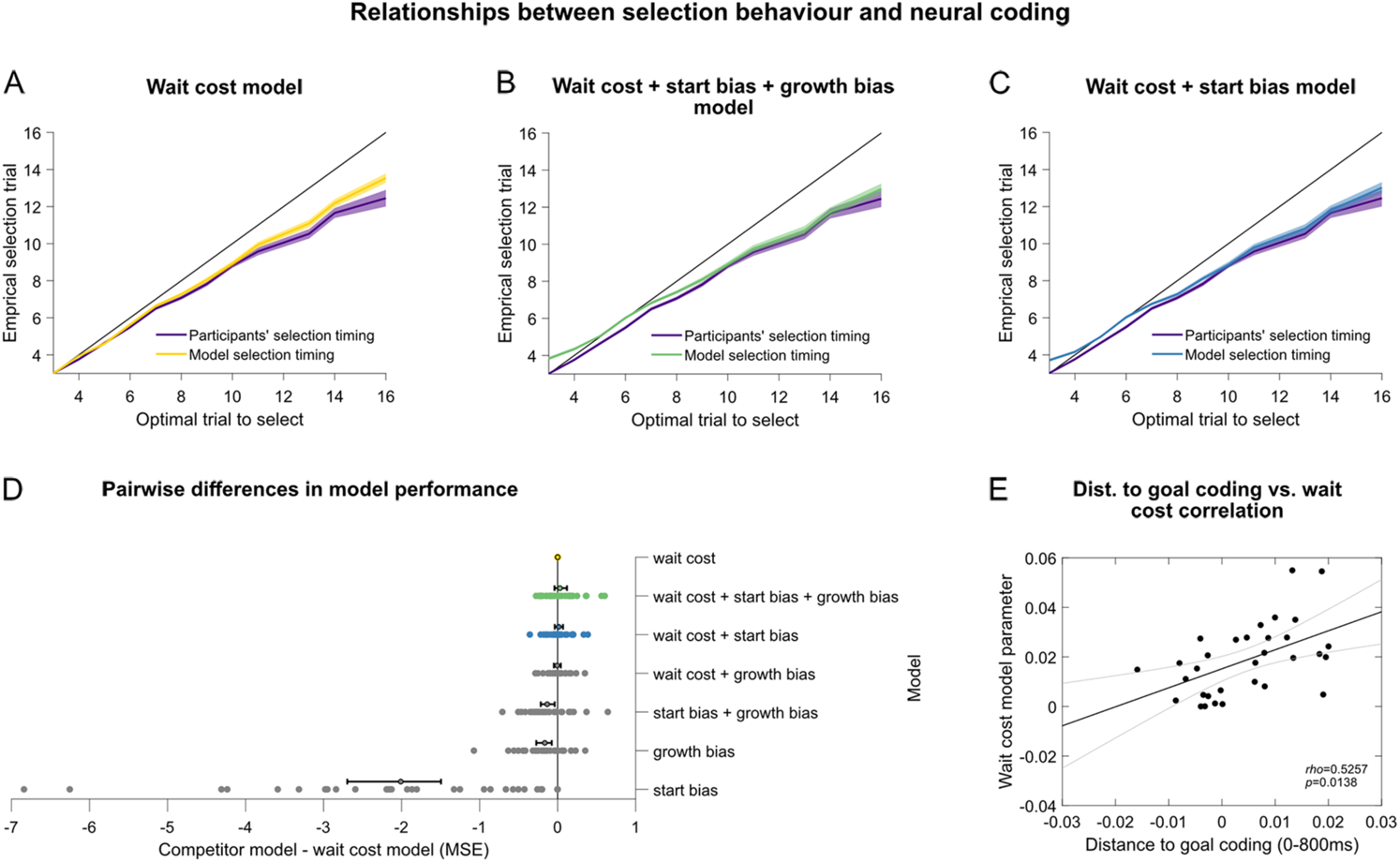
**A-C:** Top three performing models of selection timing. **A:** A cost of waiting model, which includes a reward cost proportional to number of steps waited. **B:** A model that includes a cost of waiting as well as biased representations of the starting value and growth rate. **C:** A model that includes a cost of waiting and a biased representation of the growth rate. **A-C:** Each plot shows the number of trials participants waited before selecting the option (purple line) as a function of the optimal number of trials to wait before selecting the option. The black diagonal provides a reference line for an optimal agent. The remaining coloured line shows the cross-validated performance of the model fit to participant data.

The cost of waiting model contained a single free parameter that reflected how much participants discounted an option’s reward prospect for the time that would need to be waited. A larger cost of waiting parameter meant that participants showed a stronger bias to select options early on slow trial runs. When comparing participants’ best fitting wait cost parameter with their neural data, we found a significant positive relationship between distance-to-goal encoding over the decision phase (0-800ms) in the second trial in the run and the cost of waiting parameter (Figure 4E, Spearman’s Rho=0.526, BH corrected *p*=0.014), indicating that better distance-to-goal encoding was associated with a larger bias to switch early on slow trials. No correlations were detected between the cost of waiting parameter and the neural encoding of other task variables, including the starting value on trial 1 (Spearman’s Rho=−0.200, BH corrected *p*=0.624), growth rate on trial 1 (Spearman’s Rho=−0.110, BH corrected *p*=0.549), latent reward on trial 2 (Spearman’s Rho=0.341, BH corrected *p*=0.227), distance-to-select across trials 3-6 (Spearman’s Rho=−0.228, BH corrected *p*=0.624) and select versus wait responses across trials 3-6 (Spearman’s Rho=−0.368, BH corrected *p*=0.194). To validate these correlation results in a model-free manner, we examined the relationship between the action bias (the behavioural bias to select earlier than the optimal selection trial) and neural encoding of distance-to-goal. Consistent with the results above, we found a significant relationship between the action bias and distance-to-goal encoding (Spearman’s Rho=0.471, BH corrected *p*=0.039) but not the remaining neural variables (max. Spearman’s Rho=0.373, min. BH corrected *p*=0.177).

Shading around each coloured line shows the standard error of the mean. **D:** The difference in mean squared error (MSE) between a given model and the cost of waiting model. More negative values indicate worse model performance, relative to the cost of waiting model. The y-axis shows each model being compared to the costs of waiting model. The x-axis shows the difference in MSE. Mean differences across the sample are shown with dots enclosed in a black circle and 95% confidence intervals are shown with black lines extending from the sample means. MSE differences for individual participants are shown for the top competitor models as coloured dots and the remaining competitor models as grey dots. **E:** Correlation between the average distance-to-goal coding during the decision phase (0-800ms) and the cost of waiting parameter estimates. Black lines indicate linear fits to the data and grey lines indicate 95% confidence intervals of the fits.

## Discussion

The present study examined how people decide when to select a specific option as its reward changed reliably over time, in the absence of direct sensory cues to indicate its changing value. We hypothesized that an option’s starting value and growth rate could be used to track the option’s latent reward over time, allowing the option to be selected when its reward reached a certain threshold. We assessed whether this strategy could in principle be dissociated from an alternate strategy that might lead to similar neural responses, namely determining the optimal selection time at the beginning of a trial run and waiting for this time point to occur, without continuously updating a latent reward estimate. While starting value and growth rate were strong predictors of when an option would be selected, we did not find neural evidence consistent with latent reward tracking in the present experiment. Instead, we found neural evidence that people encoded option properties and used this information to evaluate when an option would reach its maximum value, far in advance of selecting it. This distance-to-goal information was present in the data even when controlling for the option’s latent reward but not vice versa. Following the distance-to-goal signal encoded early in a trial run (on trial 2), we observed patterns of neural activity indicating how many steps into the future the option would actually be selected (on trials 3-6). These results provide evidence that even strategies that lead to very similar predictions about neural activity can be dissociated using representational similarity analysis. In the present study, people appeared to decide when to select the option in advance and monitored the time until their selection response, as opposed to actively tracking the option’s changing reward prospect.

Computational modelling further showed that selection behaviour could be captured with a temporal discounting model, which discounted an option’s maximum reward based on the time needed for its maximum to be reached. The core parameter in this model showed a positive correlation with the neural encoding of distance-to-goal information.

The present results add to recent studies examining when a specific option should be selected (Khalighinejad et al., 2020a, 2020b; Stoll et al., 2016). Research by Khalinghinejad and colleagues (2020a, 2020b) showed that recent task history is integrated with on-going sensory evidence about an option’s reward, to determine when it will be selected. Our results suggest that when on-going sensory evidence is unavailable and reward increases at a predictable rate, temporal predictions about when an option will become most valuable can be used to determine selection timing, circumventing the need to integrate changing reward information. The present results also build on findings from Stoll and colleagues (2016), who found a transient signal in the cingulate cortex encoding latent information about an option’s progress to a reward threshold. While Stoll et al.’s (2016) results raise the possibility that an option’s latent reward could be tracked over time to guide select decisions, we did not find evidence for latent reward tracking. There could be at least four explanations for this difference. First, while Stoll et al. (2016) used options with different growth rates, allowing latent reward progress and distance-to-goal to be dissociated in principle, the analysis did not explicitly dissociate the two factors. The transient signal observed in Stoll et al. (2016) could therefore reflect, in part, distance-to-goal information, signaling how soon an option would reach the reward threshold. In our particular case, the neural data was initially consistent with both kinds of timing information, requiring careful dissociation to identify the latent information being used. Future studies could take this insight into account and use dissociation procedures, such as the one in this experiment, to understand the precise mechanism guiding selection behaviour. Second, while we used a multi-stage RSA procedure to dissociate the two factors, the variables were still closely related. This could have meant that latent reward was encoded in the neural signal but could not be detected after a large amount of correlated variance, which could also be captured with distance-to-goal, was removed. To decorrelate the two variables, future designs could control changes in latent reward using a step-function, as this would allow distance-to-goal to decrease with each successive trial, while latent reward is held constant for multiple trials before increasing to the next reward step. Third, our task continued to present the starting value on each trial, which could have created interference with latent reward updating and made this variable more difficult to decode. Future designs could therefore consider showing starting value on the first trial, and presenting elapsed timesteps thereafter to demarcate changes from one trial to the next. Fourth, different environmental constraints could promote different cognitive strategies. In Stoll et al. (2016), macaques did not know how long it would take for the option to reach the reward threshold when beginning a trial run and could check the option during the run to gain information. In our study, participants had all the information needed to determine when an option would become most valuable at the start of a run, which could have encouraged the use of distance-to-goal as a proxy for changing reward values. Distance-to-goal tracking might therefore be considered the more computationally efficient solution for the current task, in which changes in reward followed a predictable trajectory during each run. One implication is that the task structure could have allowed participants to circumvent a more demanding latent reward tracking process, which might be used in more complex settings. Future studies could use the analysis approach adopted here to investigate this idea and test whether latent reward tracking might be favoured when changes in reward are less reliable. For example, options that change growth rate at unpredictable times during a trial run could make it more difficult to use distance-to-goal information, which would need to be recomputed after each change in the growth rate. The two tracking strategies could also be assessed behaviourally in future studies, by introducing probe questions at random timesteps during the run that ask participants to report the option’s current reward or the number of steps until its maximum. The latent reward account would predict that participants can report the option’s current reward with greater accuracy than the distance-to-goal. The distance-to-goal account would predict that participants can report the distance-to-goal with greater accuracy than the option’s current reward.

One interesting aspect of the results was that distance-to-goal information was detected transiently, appearing on trial 2 but not on trials 1 or 3. One speculative reason for this could be that most participants computed distance-to-goal information on trial 1 but at variable times across trial runs and across participants, making the decoding noisier. It could also be that trial 1 was noisier in general as participants settled into the run, lowering the overall decoding accuracy for distance-to-goal compared with trial 2. An alternative possibility could be that participants waited until trial 2 to compute the distance-to-goal because options in the task never reached the 500-point maximum before trial 3 and, thus, this information was not needed to make wait-select decisions on trials 1 and 2. On trial 3, distance-to-goal could have become more difficult to decode due to increasing variability in the encoding, as participants switched from thinking about the optimal selection timing to when they would actually select the option. While the distance-to-select decoding seen on trials 3-6 provides some evidence towards this, the reason distance-to-goal coding was so transient remains elusive. For example, it could also be that decoding became more difficult on trial 3 due to a general increase in recording noise further into a trial run, as participants became more distracted. Exploratory analyses, following up the primary effects, showed that the distance-to-goal signal on trial 2 was numerically stronger for options involving lower starting values. Such options had higher goal distances on average and thus stronger coding could be due to lower starting values, higher goal distances, or a combination. The distance-to-goal signal on trial 2 was also strongest in the posterior electrodes. This exploratory result might provide an interesting spatial marker of distance-to-goal information, to examine in future studies. While it is challenging to draw inferences about underlying neural sources based on M/EEG channels due to the inverse problem (Baillet & Garnero, 1997), one speculation that could be tested with more spatially specific methods would be that the posterior profile seen here reflects timing signals arising from parietal cortex. Previous studies have implicated parietal cortex as a critical region in action timing, based on findings that neuronal firing rates in the lateral intraparietal sulcus are modulated by the wait time before a motor response (Jazayeri & Shadlen, 2015). Specifically, neurons in this region gradually ramp up activity towards a fixed response threshold, with slower ramping as wait intervals become longer. These findings suggest that neurons within parietal cortex are sensitive to both the prospective wait duration and elapsed time (Jazayeri & Shadlen, 2015; see Paton & Buonomano 2018 for review), making it a possible candidate region for encoding temporal variables in the present task. Future studies could therefore use more spatially specific methods to test whether distance-to-goal information is encoded in parietal cortex, as opposed or in addition to other brain regions implicated in selecting timing, such as the basal ganglia (Khalighinejad et al., 2020a, 2020b) and dorsal anterior cingulate cortex (Stoll et al., 2016).

In addition to the neural results, we found a strong behavioural bias to select early, when options were slow to reach their maximum reward. While unanticipated, this result is consistent with findings showing humans and macaques do not wait for options to reach their full expected value before selecting them (Khalighinejad et al., 2020a, 2020b), and that the probability of making a select response in general increases over trials in sequential tasks (Baumann et al., 2020). Our computational modelling results suggest that the bias could be explained by a temporal discounting process, in which an option’s subjective value was reduced with increasing wait times (see van den Bos & McClure, 2013), rather than a biased representation of task variables that lead to inaccurate reward estimates. One aspect the modelling results do not clearly delineate is whether discounting occurred at the start of a trial run, when an option was first being evaluated, or whether it built up during the run, due to a declining ratio between the maximum reward and the cognitive effort involved in waiting longer (see Frömer et al., 2021; Kool & Botvinick, 2018; Shenhav et al. 2013; Westbrook et al., 2020; Yee & Braver, 2018). A final interesting aspect was that participants with stronger distance-to-goal encoding tended to show a stronger action bias, suggesting that those with more accurate predictions about peak reward timing selected slow options earlier. One speculative reason could be that this behaviour is beneficial in real-world environments, where decisions to select low reward rate options prematurely give decision makers more time to encounter and exploit more valuable options. This proposal is consistent with findings that human choices to abandon options are sensitive to opportunity costs (Constantino & Daw, 2015) and that people show reduced cognitive control performance when opportunity costs are high (Otto & Daw, 2019).

To conclude, we found that when option rewards followed reliable trajectories and there were no direct sensory changes to indicate changes in reward, people encoded the time when options would become most valuable in the future and monitored the number of actions until the point of selection. In contrast, we did not find evidence for independent neural coding of an option’s latent reward over time. These results suggest that, in structured environments where timing provides a useful proxy for changing reward, the human brain uses time-oriented encoding to make decisions about when to select an option. At a broader level, the approaches used in this experiment could be useful for dissociating how closely related variables impact neural activity during decision-making.

## Acknowledgements

This research was support by the Rutherford Foundation and William Georgetti Trusts of New Zealand (S.H.), the Alexander von Humboldt Foundation (S.H.), the Wellcome Trust (award 201409Z/16/Z to N.E.M), University College Oxford (N.E.M.), the Biotechnology and Biological Sciences Research Council (award BB/M010732/1 to M.G.S.), the James S. McDonnell Foundation (award 220020405 to M.G.S.), and the NIHR Oxford Health Biomedical Research Centre. The Wellcome Centre for Integrative Neuroimaging is supported by core funding from the Wellcome Trust (203139/Z/16/Z). The authors would like to thank Keno Jüchems and Joshua Calder-Travis for comments on previous versions of the manuscript, Fabrice Luyckx for comments on the figures, as well as Nils Kolling, Laurence Hunt, Roshan Cools and Matthew Rushworth for helpful discussions.

## Author Contributions

S.H, M.G.S and N.E.M designed the study. S.H acquired the data. S.H and N.E.M. analysed the data with input from M.G.S. S.H wrote the original draft. S.H, N.E.M and M.G.S reviewed and edited the manuscript.

## Diversity Statement

Here we report the predicted gender and ethnicity of authors in our reference list (Ambekar et al., 2009; Bertolero et al., 2020; Dworkin et al., 2020; Sood et al., 2018; Zhou et al., 2020). First, we obtained the predicted gender of the first and last author of each reference by using databases that store the probability of a first name being carried by a woman. By this measure (and excluding self-citations to the first and last authors of our current paper), our references contain 2.5% woman(first)/woman(last), 12.5% man/woman, 25% woman/man, and 60% man/man. This method is limited in that a) names, pronouns, and social media profiles used to construct the databases may not, in every case, be indicative of gender identity and b) it cannot account for intersex, non-binary, or transgender people. Second, we obtained the predicted ethnicity of the first and last author of each reference by databases that store the probability of a first and last name being carried by an author of colour. By this measure (and excluding self-citations), our references contain 4.29% author of colour(first)/author of colour(last), 13.76% white author/author of colour, 23.11% author of colour/white author, and 58.84% white author/white author. This method is limited in that a) names and Florida Voter Data to make the predictions may not be indicative of ethnicity, and b) it cannot account for Indigenous and mixed-race authors, or those who may face differential biases due to the ambiguous ethnicisation of their names.

## Notes

Conflict of interest: The authors declare no competing financial interests.

### Competing Interest Statement

The authors have declared no competing interest.

